# NT3/TrkC pathway modulates the expression of UCP-1 and adipocyte size in human and murine adipose tissue

**DOI:** 10.1101/2020.07.24.216374

**Authors:** María Bové, Fermi Monto, Paloma Guillem-Llobat, M Dolores Ivorra, M Antonia Noguera, Andrea Zambrano, Ma Salome Sirerol-Piquer, Ana Cristina Requena, Mauricio García-Alonso, Teresa Tejerina, José T. Real, Isabel Fariñas, Pilar D’Ocon

**Author notes:** **Corresponding author:** Pilar D’Ocon, Departamento de Farmacología, Facultad de Farmacia and Estructura de Recerca Interdisciplinar en Biotecnologia i Biomedicina (ERI BIOTECMED), Universidad de Valencia, Avda. Vicent Andres Estelles s/n, Burjassot, Valencia, 46100, Spain.

## Abstract

NT3, through activation of its tropomyosin-related kinase receptor C (TrkC), modulates neuronal survival and neural stem cell differentiation. It is widely distributed in peripheral tissues (specially vessels and pancreas) and this ubiquitous pattern suggests a role for NT3, outside the nervous system and related to metabolic functions. The presence of the NT3/TrkC pathway in the adipose tissue (AT) has never been investigated. Present work studies in human and murine adipose tissue (AT) the presence of elements of the NT3/TrkC pathway and its role on lipolysis and adipocyte differentiation. qRT-PCR and immunoblot indicate that NT3 was present in human retroperitoneal AT and decreases with age. NT3 was also present in rat isolated adipocytes and retroperitoneal, interscapular, perivascular and perirenal AT. Histological analysis evidences that NT3 was mainly present in vessels irrigating AT close associated to sympathetic fibers. Similar mRNA levels of TrkC and β-adrenoceptors were found in all ATs assayed and in isolated adipocytes. NT3, through TrkC activation, exert a mild effect in lipolysis. Addition of NT3 during the differentiation process of human pre-adipocytes resulted in smaller adipocytes and increased uncoupling protein-1 (UCP-1) without changes in β-adrenoceptors. Similarly, transgenic mice with reduced expression of NT3 (Ntf3 knock-in lacZ reporter mice) or lacking endothelial NT3 expression (Ntf3flox1/flox2;Tie2-Cre+/0) displayed enlarged white and brown adipocytes and lower UCP-1 expression.

**Conclusions:** NT3, mainly released by blood vessels, activates TrkC and regulates adipocyte differentiation and browning. Disruption of NT3/TrkC signaling conducts to hypertrophied white and brown adipocytes with reduced expression of the thermogenesis marker UCP-1

## 1. INTRODUCTION

The neurotrophin family of proteins is classically known for the effects of its members on neuronal survival. All NTs or their (pro)NT forms bind to the low-affinity p75 neurotrophin receptor (p75NTR), a member of the tumor necrosis factor receptor superfamily, whereas each mature NT interacts with high-affinity with a specific membrane tyrosine kinase receptor of the tropomyosin-related kinase receptor family (Trk). Nerve growth factor (NGF) preferentially binds TrkA, nerve growth factor (NGF), brain-derived neurotrophic factor (BDNF) and neurotrophin 4/5 preferentially bind TrkB, and neurotrophin-3 (NT3) preferentially binds TrkC, although it can also signal through TrkA and TrkB [1,2].

NGF and BDNF mediate not only neurotrophic [3], but also metabotropic effects, having been implicated in obesity, diabetes mellitus and metabolic syndrome [4]. There is evidence indicating that NGF and BDNF are produced by the adipose tissue (AT) [5,6,7], being designated as “metabokines” [8,9] due to their role as modulators of energetic metabolism.

The best known activity of NT3 is related to its ability to modulate neuronal survival of sensory and adrenergic neurons during embryonic development [10–15]. In the developing central nervous system, NT3 is involved in the growth of connections and synapse formation [16, 17] but its levels dramatically decrease with brain maturation [18]. In contrast, NT3 is widely distributed in peripheral tissues of adult animals and it has been detected in vessels, pancreas, spleen, liver, skin, gut, skeletal muscle, lung and heart [19–24]. This ubiquitous pattern suggests that NT3 could play an essential role outside the nervous system, especially in pancreas [22] and vessels [25] where its level of expression is elevated, and opens the possibility for a role of this neurotrophin also in the control of metabolic processes. However, the presence of molecular elements of the NT3/TrkC pathway and their activity in the different types of AT has not been investigated so far.

Only indirect studies analyzed the impact of central depletion of NT3 or TrkC overexpression on mice body weight, but changes were not evident [3]. In humans, Bullo et al. [26] quantified plasma levels of NT3 in women with different degrees of adiposity, obesity and metabolic syndrome. NT3 levels tend to be higher in morbidly obese patients, and seems to be related to lipid profile since NT3 levels negatively correlates with the concentration of total cholesterol and low-density lipoprotein (LDL)-cholesterol, whereas lower NT3 levels were found in patients with hypertriglyceridemia. Different results were observed by Popa-Wagner et al. [27] working with a group of alcoholic men. These authors also described elevated levels of NT3 in serum of obese patients, being positively associated with body mass index (BMI), but in this case, a positive correlation was found between plasmatic NT3 levels and LDL-cholesterol.

Hence, the objective of this research was to investigate the expression and function of NT3/TrkC pathway in human and rodent AT, as well as its relationship to the β-adrenergic system, the main modulator of AT function. Interestingly, we have found the presence of elements of the NT3/TrkC pathway in white and brown AT (WAT and BAT, respectively), their relationship to lipid metabolism, and the direct involvement of NT3 in adipocyte differentiation and uncoupling protein-1 (UCP-1) expression, the major determinant of non-shivering thermogenesis and a characteristic marker of brown and brite adipocytes [28]. Together, our data point at NT3/TrkC as new potential targets to regulate adipocyte differentiation and browning

## 2. METHODS

### 2.1. Patient characteristics

This research conforms with the principles outlined in the Declaration of Helsinki and was performed with the approval of the Ethic Committee of the Hospital Clínico Universitario San Carlos, Madrid, Spain with registry number 19/011. Patients (n = 28) were consecutively recruited at the General Surgery Service of the Hospital Clínico Universitario San Carlos, Madrid (Spain), from those undergoing abdominal surgery by colon neoplasia (n = 23) or other pathologies (n = 5). Subjects were included if they were free of overt inflammatory or infectious diseases. The presence of diabetes or hypertension was confirmed when a previous diagnosis. Height and weight were measured without shoes and with light clothes, and BMI was calculated as weight (kg)/height (m)^2^. Venous blood samples were taken at 8.00 a.m. under overnight fasting conditions. Plasma was immediately separated by centrifugation and glucose, triglycerides, and lipid profile were assessed by the hospital’s routine chemistry laboratory. Abdominal retroperitoneal white AT, normally discarded, was collected during the surgical procedures, kept in RPMI 1640 (Gibco) medium at 4 °C, and frozen at −80 °C immediately until used. Only tissues devoid of any obvious lesions were used. Clinical and anthropometric characteristics of patients included in the study were summarized in Table 1.

**Table 1.** Demographical characteristics and biochemical profile of the subjects included in the study.

|  |  |
| --- | --- |
| **Sex, m/w** | 15/13 |
| **Age, years** | 73.5 ± 2.1 [ 41 - 90 ] |
| **BMI. kg/m<sup>2</sup>** | 25.5 ± 0.7 [ 19 - 32 ] |
| **Glucose, mg/dL** | 97.0 ± 3.7 [ 64 – 161 ] |
| **Total cholesterol, mg/dL** | 182.4 ± 6.5 [ 128 - 245 ] |
| **HDL-cholesterol, mg/dL** | 53.8 ± 3.1 [ 31 - 93 ] |
| **LDL-cholesterol, mg/dL** | 99.6 ± 5.1 [ 63 - 147 ] |
| **Triglycerides, mg/dL** | 114.3 ± 12.3 [ 55 - 286 ] |
| **HTA, m/w** | 9/6 |
| **Diabetes, m/w** | 4/3 |
| **Smoking, m/w** | 2/1 |
| **Sedentary lifestyle, m/w** | 4/6 |

m=men; w = women

### 2.2. Animals and *in vivo* procedures

*Ntf3^+/lacZneo^* mice [10] bred in CD1 genetic background for more than ten generations [24], and *Ntf3^flox1/flox2^* and *Tie2-Cre^+/0^* mice obtained from The Jackson Laboratory (U. S.) were genotyped as previously described [10, 29, 30]. Mice of the different strains and Wistar rats were bred in our in-house animal facility. All animals were gender group-housed with standard pelleted food and tap water *ad libitum*. Animal handling and all experimental procedures comply with the ARRIVE guidelines and were carried out according to European Union 2010/63/UE and Spanish RD-53/2013 guidelines, following protocols approved by the institutional ethics committee of the University of Valencia. Animals were deeply anaesthetized with isoflurane. To determine glycemia, blood samples were collected from the saphena vein by skilled personnel. After this, animals were killed by transcardial blood withdrawal and blood was placed in heparinized tubes and centrifuged at 1500*g* at room temperature for 30 min in an Eppendorf Centrifuge 5804-R (Hamburg, Germany) to obtain plasma which was immediately frozen at −80 °C prior to analysis. Cerebral cortex, left ventricle, retroperitoneal white AT (WAT), interscapular brown AT (BAT), perivascular AT surrounding aorta (VAT) and AT surrounding kidney (perirenal, KAT) were removed. Following dissection, tissues were cleaned with phosphate buffered saline solution (PBS), frozen in liquid nitrogen, and stored at –80 °C or placed in PBS for immediate use. The glycemia was determined in a glucometer (Contour XT ^®^ Bayer). Lipid profile was determined in plasma using an autoanalyzer (Gernonstar^®^, Ansasia, Bombay, India).

### 2.3. Histological analysis

Freshly isolated WAT and BAT tissues from male mice were immersion-fixed in 4% paraformaldehyde (PFA) in PBS at pH 7.4 overnight, dehydrated, embedded in paraffin and sectioned into 5 μm-sections. Samples were deparaffinized, rehydrated and stained with haematoxylin and eosin for the evaluation of adipocyte size. Digital images of AT sections were captured using a light microscope (Leica DM IL LED, Germany) at 10X magnification. For each experimental group, five sections *per* mouse were stained and the cell areas of 20 adipocytes *per* section were measured using image analysis software (ImageJ; U. S. National Institutes of Health, Bethesda, Maryland, USA). Some fixed samples were reacted with the X-gal histochemistry for detection of β-galactosidase (β-gal) expression. The samples were incubated in PBS with 2 mM MgCl_2_, 5 mM K_3_Fe(CN)_6_, 5 mM K_4_Fe(CN)_6_, 0.01% sodium deoxycholate and 0.02% NP-40 and 1 mg/ml X-Gal for 24 h and mounted in glycerol. Some mice were deeply anesthetized under veterinarian supervision and injected transcardially with 100 μL of 0.8 mg/ml FITC-conjugated IB4 lectin in saline solution for 1 min before extensive perfusion with saline solution. After euthanasia fat samples were immersion-fixed in 4% PFA, washed three times in PBS containing 2% Triton X-100 for 15 min each, blocked for 2 h in 10% FBS and 2% Triton X-100 in PBS, then incubated with primary antibodies: SMA1 (1:200, Abcam Ab96361), β-gal (1:500 Abcam Ab7817) and TH (1:500, Novus NB300-109). Detection was performed with fluorescent secondary antibodies: Cy3 donkey-α-chicken from Jackson ImmunoResearch, Alexa Fluor^®^ 546 donkey-α-rabbit and Alexa Fluor^®^ 647 donkey-α-mouse from Life Technologies, at 1:800 in the same blocking solution. Fluorescent samples were mounted with Fluorsave (Calbiochem) and analyzed with an Olympus FV10 confocal microscope.

### 2.4. Adipocyte isolation

WAT was removed and minced with scissors in Krebs-Ringer containing 15 mM sodium bicarbonate, 10 mM HEPES, and bovine serum albumin 3.5% (w/v), pH 7.4. The tissue was digested for 35 to 45 min at 37°C with 1.5 mg/ml collagenase. Isolated fat cells were washed three times in a large amount (around 20 ml) of the same buffer without collagenase. After washed, the floating fat cells were frozen in liquid nitrogen, and stored at –80 °C or placed in around 10-fold their volume of the same buffer for immediate use

### 2.5. RNA isolation and expression analysis

To obtain total RNA, frozen samples were treated as previously described [31]. 500 ng of RNA were reverse-transcribed into cDNA using oligo(dT)16 and ImProm-II TM Reverse Transcriptase (Promega Corp., Madison, USA). cDNA products were amplified on a GeneAmp 7500 Fast System (Applied Biosystems, Carlsbad, CA, USA), using TaqMan^®^ Gene Expression Assay Mix (Thermo) as previously described [32]. The specific primer-probes were: *NTF3* (Hs01548350_m1); *NTRK3* (Hs00176797_m1); *ADRB1* (Hs00265096_s1); *ADRB2* (Hs 00240532_s1); *ADRB3* (Hs 00609046_m1) and *GAPDH* (Hs99999905_m1) for human samples. *Ntf3* (Rn00579280_m1)*; Ntrk3* (Rn00570389_m1); *Adrb1* (Rn008245936_s1); *Adrb2* (Rn00560650_s1); *Adrb3* (Rn00565393_m1) and *Gapdh* (Rn99999916_s1) for rat samples. *Ntf3* (Mm01182924_m1)*; Ntrk3* (Mn00456222_ m1); *Adrb1* (Mm01265414_s1); *Adrb2* (Mm02524224_s1); *Adrb3* (Mm00442669_m1) and *Gapdh* (Mm99999915_g1) for mouse samples. cDNA was amplified following the manufacture’s conditions. The targets and reference were amplified in parallel reactions. The Ct values obtained for each gene were referenced to *GAPDH* (human) or *Gapdh* (rodents), and converted into the linear form using the term 2^-ΔCt^ as a value directly proportional to the mRNA copy number.

### 2.6. Immunoblot

Frozen tissues were pulverized and a protein extract was obtained and stored as previously described [33]. 30 μg of frozen protein extracts were incubated with the SDS-sample buffer (2% SDS, 60 mM Tris-HCl buffer pH 6.8, 5% β-mercaptoethanol, 0.01% bromophenol blue and 10% glycerol), separated on 10-15% SDS-polyacrylamide gels and transferred to polyvinylidene fluoride (PVDF) membranes for 2 h at 340 mA, using a liquid Mini Trans-Blot^®^ Electrophoretic Transfer Cell system (Bio-Rad Laboratories, Inc.). Membranes were blocked in 5% bovine serum albumin in Tris buffered saline (TBS) containing 0.1% Tween 20 (TBST) for 1 h at room temperature with gentle agitation. Membranes were incubated overnight at 4 °C with the indicated primary antibodies diluted in blocking solution: NT3 (1:1000, Abcam ab65804), TrkC (1:750, Cell Signaling Technologies C44H5), UCP1 (1:1000, Sigma-Aldrich U6382) and Gapdh(1:10000, Sigma-Aldrich G9545). Membranes were then washed three times, incubated with ECL^™^ peroxidase labelled goat anti-rabbit IgG (Cell Signaling Technologies 7074S) (1:2500) for 50 min at room temperature, and washed extensively before developing by incubation with the ECL^®^ western blotting detection reagent (Amersham Biosciences). Membranes were immediately documented and quantified with an Autochemi™ BioImaging System using the Labworks 4.6 capture software (Ultra-Violet Products Ltd., Cambridge, UK).

### 2.7. Lipolytic Activity in Isolated Adipocytes

Glycerol concentration was determined as an index of lipolytic activity, and was enzymatically measured using glycerokinase and NAD-dependent glycerophosphate dehydrogenase as previously described [34]. A 400 μl aliquot of the well-stirred cell suspension of isolated adipocytes was immediately distributed into plastic incubation vials containing 4 μl of drug dilutions at 100× the final concentration to be tested. Isolated adipocytes were incubated in Krebs-Ringer with sodium bicarbonate (15 mM), HEPES (10 mM), and bovine serum albumin (3.5% w/v), or in the same buffer with NT3 (32 or 96 ng mL^-1^), isoprenaline (0.1, 1 or 10 μM) and 0.2 μM K252a. After a 90-min incubation, at 37°C under constant shaking and oxygenation withO2-CO2(95:5), the vials were placed in icewater for 5 min and aliquots (60 μl) of the infranatant medium were transferred to a new tube with a solution containing adenosine 5’ triphosphate (ATP, 1.5 mM, Sigma-Aldrich), β-Nicotinamide adenine dinucleotide hydrate (NAD, 0.5mM, Sigma-Aldrich), glycerophosphate dehydrogenase (GDH, 8500 units L^-1^, Merck), glycerol kinase (≥357 units L^-1^, Sigma), glycine (0.2M, Sigma), hydrazine hydrate solution (1M, Sigma) and CaCl2 2mM. The data were normalized to protein concentration and genomic DNA content. The protein concentration was determined by the Bradford reagent (Sigma-Aldrich). For each experiment, glycerol production in presence of the different drugs was calculated as a percentage *vs* glycerol determined in the vial containing adipocytes incubated with buffer but in absence of any drug (control).

### 2.8. Adipocyte differentiation

Human pre-adipocytes were purchased from Promocell (C-12732). For adipocyte differentiation, pre-adipocytes at passage 3 were cultured in preadipocyte differentiation medium (Promocell) at 37 °C in 5% CO2, following manufacturer’s conditions, Cells were differentiated from human preadipocytes to mature adipocytes in absence or in presence of different stimulus: NT3 (32 ng mL^-1^), K252a (0.2 μM) and NT3 + K252a that were added during the whole differentiation procedure. A total of 5 different experiments were performed in duplicate. Individual adipocyte size or lipid droplets diameter were determined in 6 fields by plate, and 10 cells and 25 lipid droplets were quantitated by field, using ImageJ (U.S. National Institutes of Health, Bethesda, Maryland, USA)

### 2.9. Statistical Analysis

Data were expressed as mean ± standard error of the mean. A normality test, followed by one-way analysis of variance (ANOVA). If values passed normality test, Newman-Keuls or Student’s *t* test were performed. If values do not exhibit a normal distribution, Kruskal-Wallis test followed by Dunns test were performed. To establish associations between variables parametric or not parametric, we calculated Pearson’s or Spearman’s correlation, respectively. Graph Pad Prism 4 Software was used. A probability value of *P* < 0.05 was considered significant.

## 3. RESULTS

### 3.1. NT3, TrkC and the three subtypes of β-adrenoceptors are present in human and rat adipose tissues and changes with age

As a first approach to the potential function of TrkC-mediated NT3 signaling on lipid metabolism, we analyzed their mRNA and protein levels in hAT obtained by abdominal surgery. In addition, genic expression of β1, β2- and β3-adrenoceptors (*ADRB1, ADRB2* and *ADRB3*, respectively), the main regulators of AT function, was also determined and compared to NT3 and TrkC levels. Analysis by qRT-PCR (Figure 1A), and immunoblot (Figure 1B), indicated that *NTF3* and *NTRK3* are expressed in hAT as well as *ADRB1, ADRB2* and *ADRB3* (Figure 1A). Among the membrane receptors, *ADRB2* was the most highly expressed in hAT, whereas similar levels of *ADRB1, ADRB3* and *NTRK3* were found in this tissue (Figure 1A). In spite of the lower proportion of *ADRB3* and *ADRB1*, an inverse correlation exists between *ADRB3* expression and BMI (*Spearman r* = −0.3820, *P* = 0.0448) or age (*Spearman r* = −0.3835, *P* = 0.0483). An inverse correlation was also found between *NTF3* expression in hAT and age (Spearman r = −0.4293, P = 0.0226), but no significant correlation existed between *NTF3* and *NTRK3* expression with any of the other clinical variables summarized in Table 1. Finally, mRNA levels of the five genes did not revealed changes related to gender, hypertension, diabetes, neoplasia or smoking habit (data not shown), but a significantly lower expression of *ADRB2* was found in patients with a sedentary *vs* active lifestyle (1260 ± 170, n = 10, *vs* 2775 ± 476, n = 18, *P* = 0.0292, Student’s *t* test).

**Figure 1.**
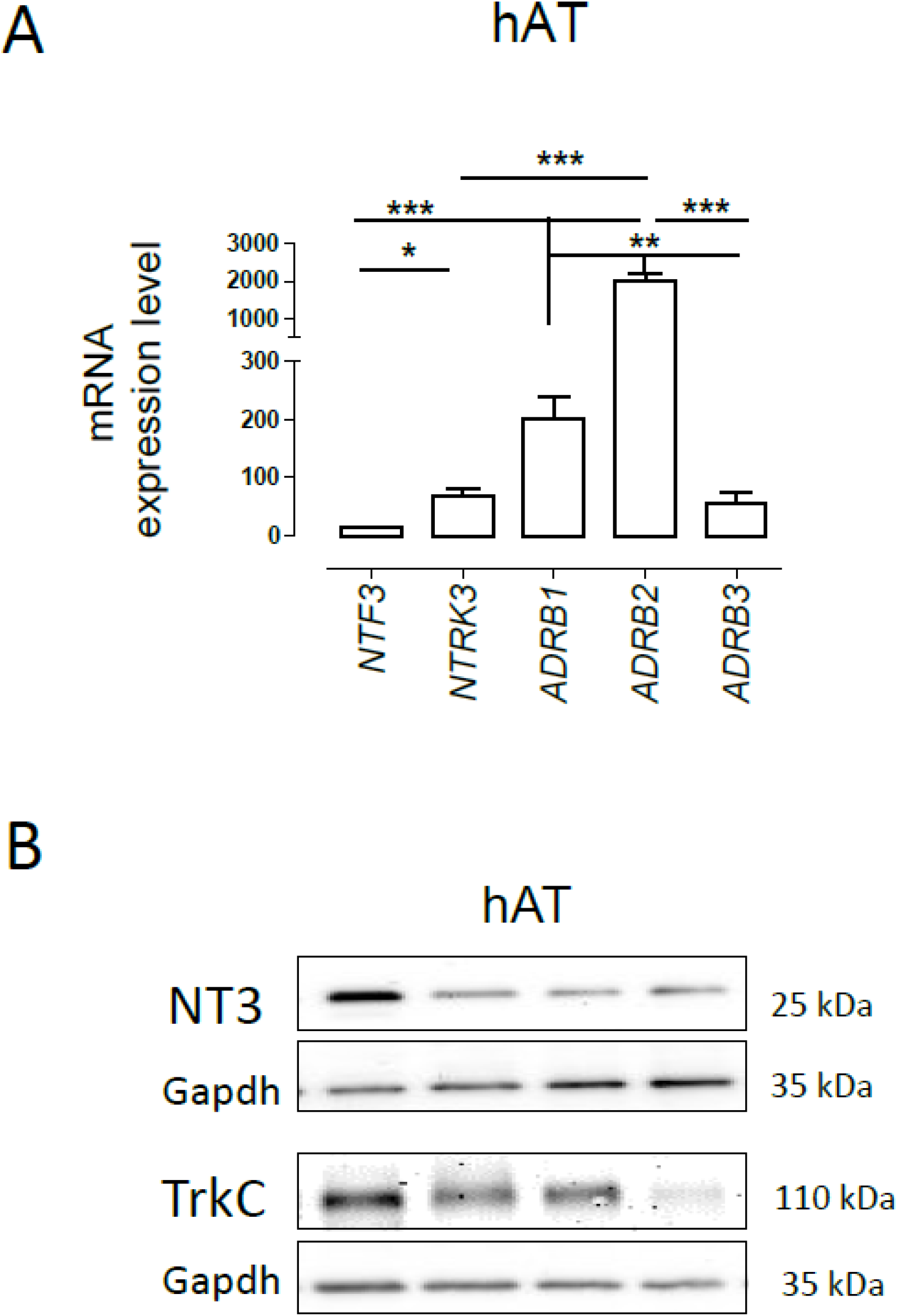
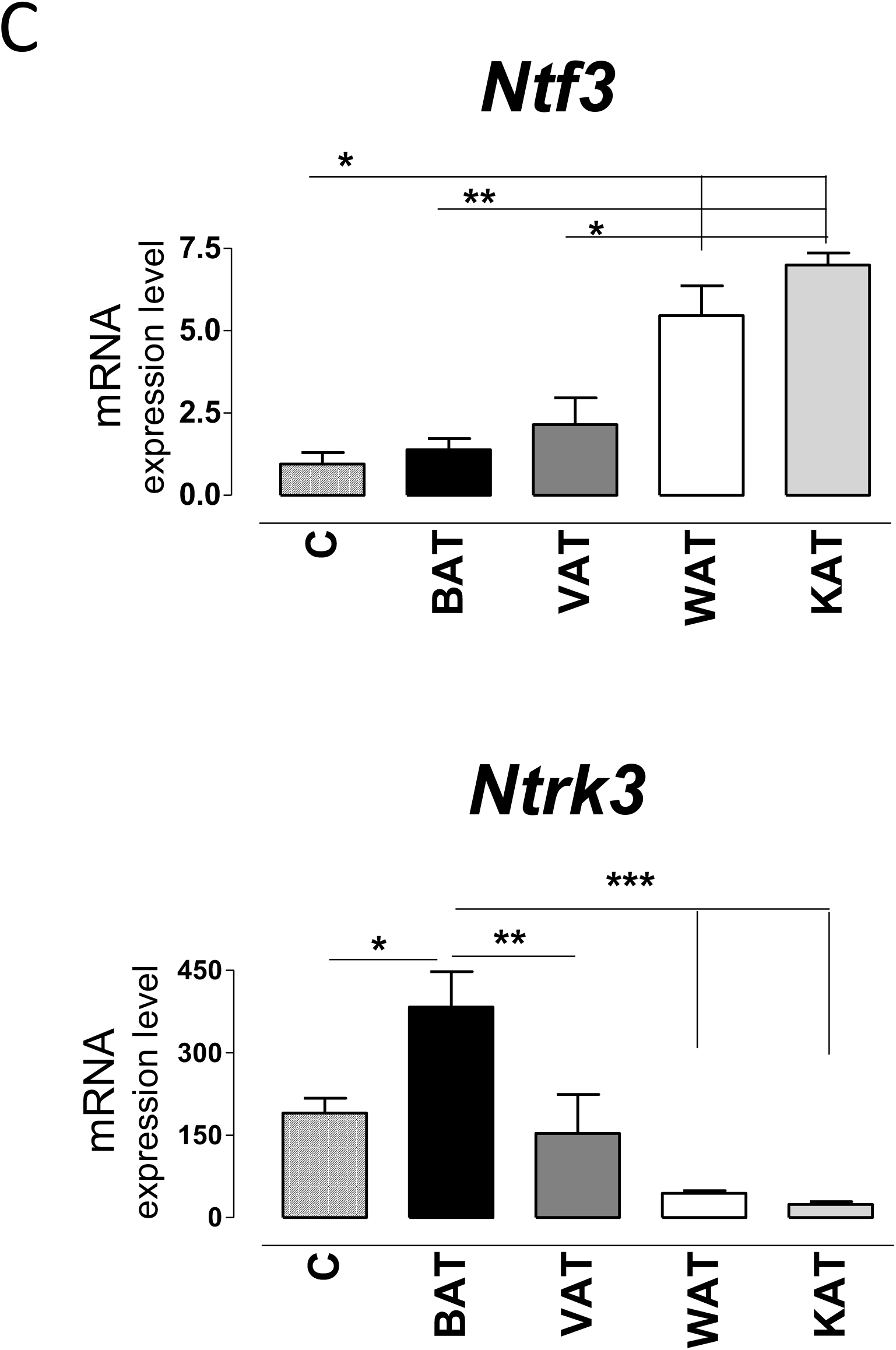
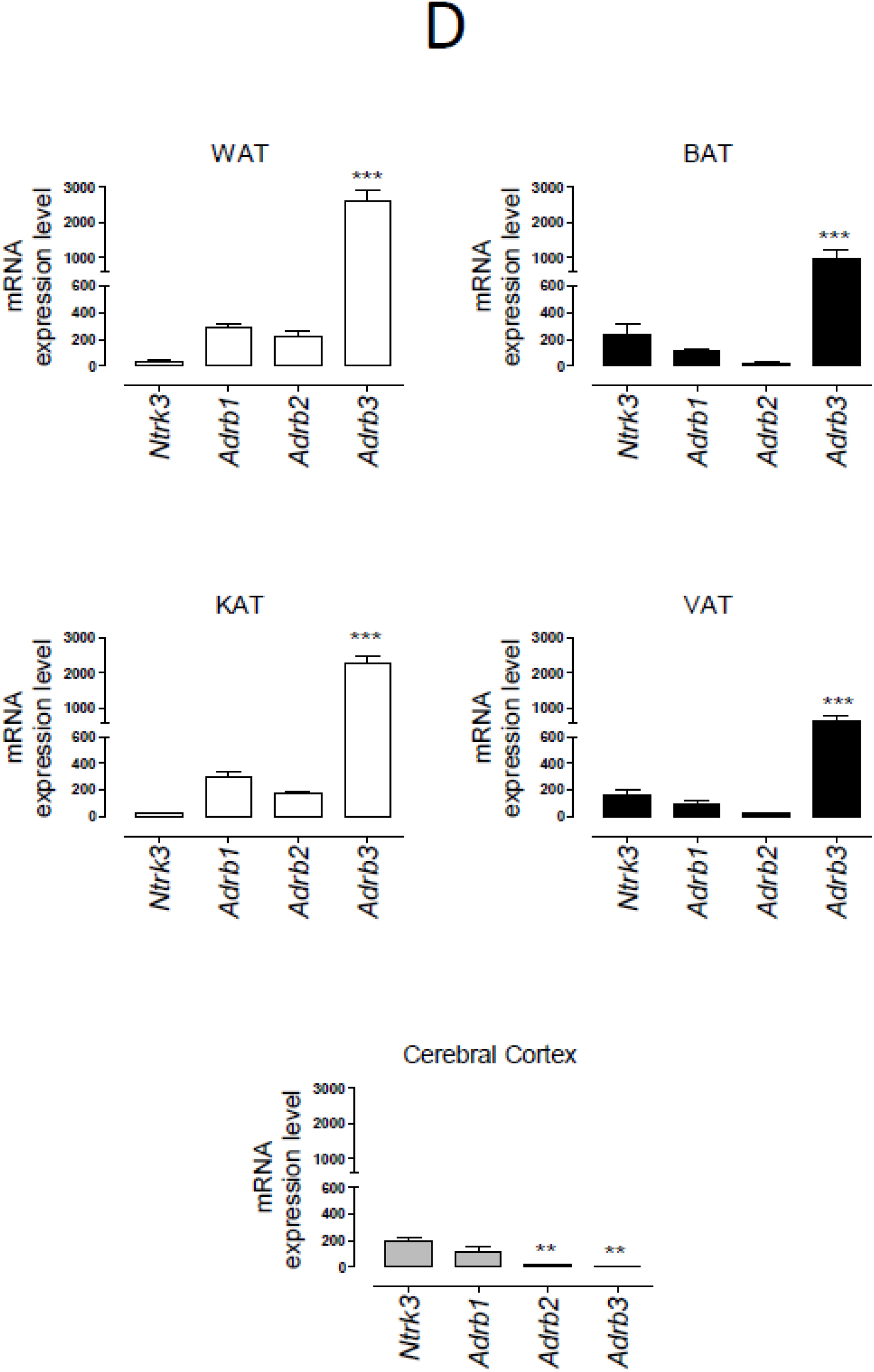
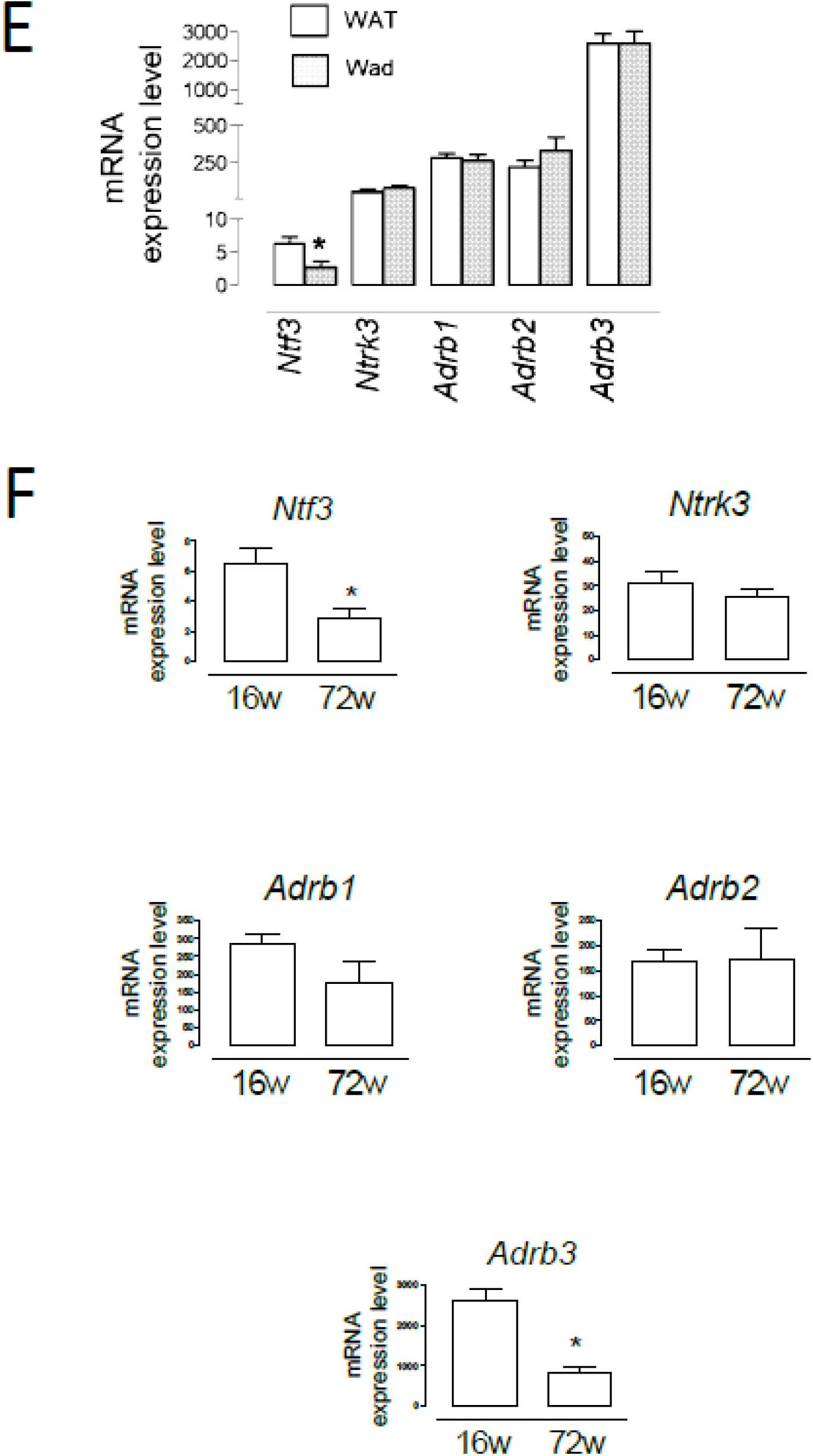
Expression of NT3 and TrkC in adipose tissues. **(A)** mRNA levels of NT3 (*NTF3*), TrkC (*NTRK3*) and the three β-adrenoceptor subtypes (*ADRB1, ADRB2, ADRB3*) in human retroperitoneal adipose tissue (hAT, n = 28) * *P* < 0.05, ** *P* < 0.01, *** *P* < 0.001, Kruskal-Wallis test followed by Dunns multiple comparison test **(B)** representative immunoblots of NT3 and TrkC. **(C)** Expression of *Ntf3* and *Ntrk3* in different tissues from Wistar rat: cerebral cortex (C, n =3), interscapular adipose tissue (BAT, n = 6), perivascular adipose tissue (VAT, n = 5), abdominal adipose tissue (WAT, n = 15) and perirenal adipose tissue (KAT, n = 5), * *P* < 0.05 ** *P* < 0.01, *** *P* < 0.001 *vs* WAT and KAT (*Ntf3*) or indicated genes (*Ntrk3*), one-way ANOVA followed by Newman-Keuls test **(D)** Comparative representation of the expression of the four membrane receptors assayed (*Ntrk3, Adrb1, Adrb2 and Adrb3*) in different tissues (WAT, BAT, KAT, VAT and cerebral cortex), * *P* < 0.05 *vs* the other genes, one-way ANOVA followed by Newman-Keuls test. **(E)** Expression of *Ntf3, Ntrk3, Adrb1, Adrb2 and Adrb3* in WAT and adipocytes isolated from WAT (Wad) (n = 8) * *P* < 0.05, Student’s *t* test *vs* WAT. **(F)** Expression of *Ntf3, Ntrk3, Adrb1, Adrb2 and Adrb3* in WAT isolated from young (16 weeks old, n = 14) and aged (72 weeks old, n = 5) Wistar rats, * *P* < 0.05, Student’s *t* test. Values of mRNA in bar diagrams are expressed as 2^-ΔCt^x10^4^ using *Gapdh* as a housekeeping gene and represent the mean ± SEM.

Expression of *Ntf3* and *Ntrk3* was also observed in the retroperitoneal WAT, the interescapular BAT, the perivascular AT (VAT) and the perirenal AT (KAT) obtained from male Wistar rats. Levels of *Ntf3* and *Ntrk3* in the different fat samples were in all similar or above those detected in extracts of cerebral cortex (C), a territory where *NTf3* and *Ntrk3* expression had been previously studied [22, 25]. It is remarkable that *Ntrk3* expression in BAT and VAT was similar to that found in cerebral cortex (Figure 1C). The expression of the three β-adrenoceptors (*Adrb1, Adrb2* and *Adrb3*) is also shown (Figure 1D). As occurred in human samples, *Ntrk3* was expressed in the rat ATs at levels comparable to β1 and β2-adrenoceptors (Figure 1D). Unlike the observation in hAT, the predominant β-adrenoceptor in rat AT is the subtype β3. Figure 1D shows the existence of a common pattern of expression, in WAT and KAT vs BAT and VAT, for the four genes of membrane receptors assayed. In adipocytes freshly isolated from rat WAT (Wad), *Ntf3* and *Ntrk3* were also detected. No significant differences were observed in the expression of the β-adrenoceptors between the whole tissue (WAT) and the isolated adipocytes (Wad). Interestingly, mRNA levels of *Ntf3* were lower in Wad (Figure 1E), suggesting that NT3 could be mainly expressed in other cell types than in adipocytes. However, *Ntrk3* expression in adipocytes was equivalent, or even tend to be higher, than that in the entire AT, supporting a functional role for TrkC in these cells.

An inverse correlation has been found between *NTF3* expression in hAT and age. In order to analyze the consequences of ageing on the NT3/TrkC pathway in rats, we compared the mRNA levels of the five genes assayed in WAT and BAT obtained from young (16 weeks old) and old (72 weeks old) Wistar rats (Figure 1F). As in hAT, a reduction in the expression of *Ntf3* and *Adrb3* was observed in WAT from aged animals, whereas *Ntrk3, Adrb1* and *Adrb2* expression remains constant. No significant changes with ageing were observed in rat BAT (results not shown, n= 4-6).

### 3.2. NT3 is expressed in adipose tissue-irrigating blood vessels in proximity to sympathetic innervating fibers

We next took advantage of the presence of a lacZ reporter gene in the targeting construct that was used for the generation of the *Ntf3^+lacZneo^* mouse strain in order to visualize the sites of NT3 expression in AT. We reacted abdominal fat pads from *Ntf3*^+/lacZneo^ young adult mice for β-gal detection and found an intense staining consistent with blood vessels (Figure 2A). We next detected the reporter enzyme with specific antibodies to β-gal (no signal in wild-type tissue) and to smooth muscle actin (SMA1) in AT from animals which had been transcardially injected prior to euthanasia with FITC-labeled lectin IB4 to specifically decorate the luminal side of endothelial cells. We observed again that β-gal expression was mainly present in blood vessels (Figure 2B). Due to the role of sympathetic innervation in AT function, we also stained the fat pads for the detection of sympathetic fibers with antibodies to tyrosine hydroxylase (TH), with either immunoperoxidase detection (Figure 2C) or immunofluorescence in lectin-injected samples (Figure 2D). The proximity of the innervating fibers to vessels, the main source of NT3, could suggest indirect neuronal effects of NT3 in AT in addition to a possible direct action in adipocytes.

**Figure 2.**
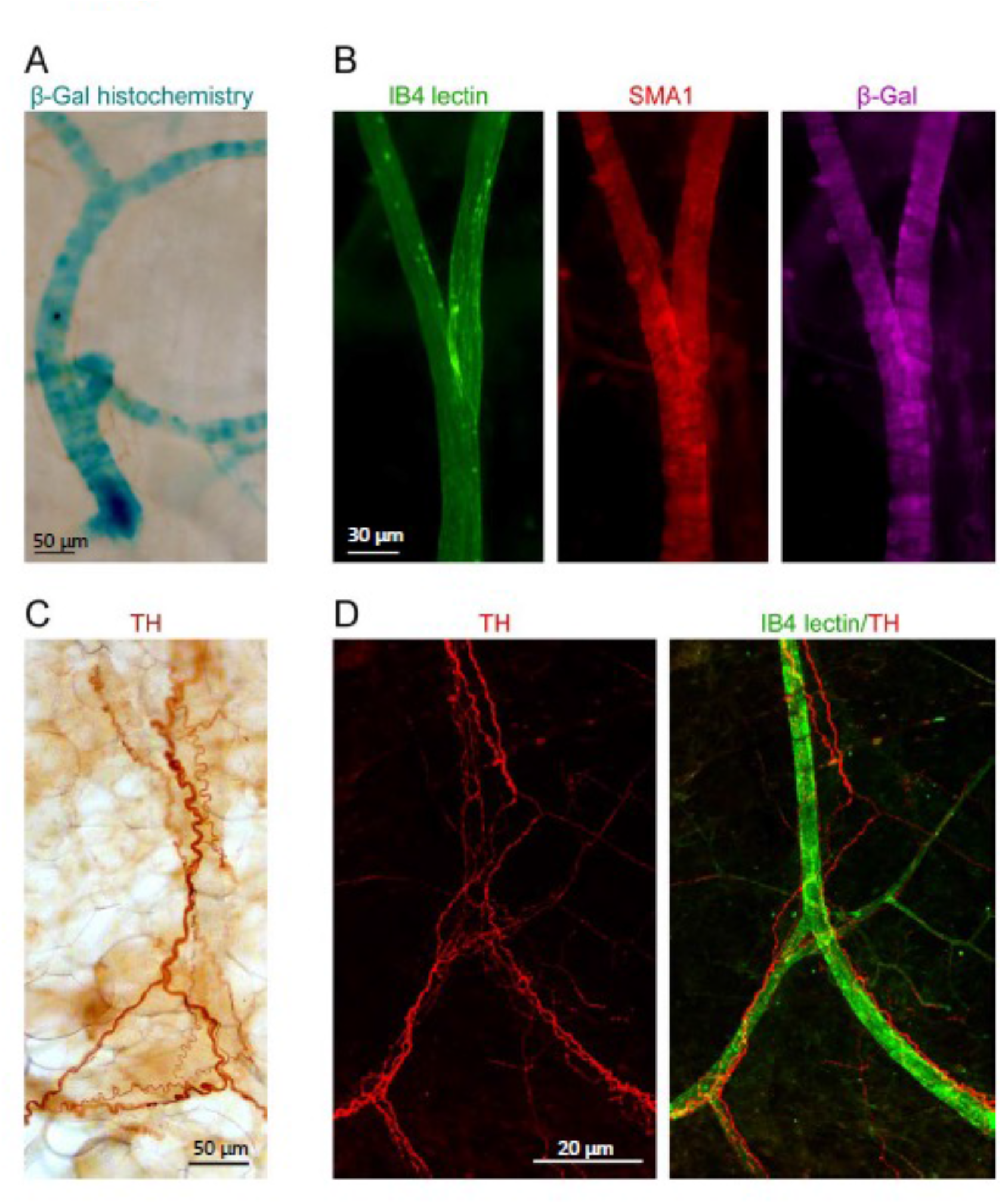
Blood vessels irrigating WAT express NT3 and are in close association with sympathetic fibers. **(A)** β-galactosidase (β-gal) histochemistry in WAT from *NTF3^+/lacZneo^* mouse showed a staining pattern consistent with blood vessels. **(B)** Immunoflorescence against SMA1 (red) and β-gal (purple) in mice transcardially injected with FITC-labeled lectin IB4 (green). Specific staining of β-gal was observed in blood vessels. **(C)** Immunoperoxidase detection of tyrosine hydroxylase (TH) and **(D)** immunoflurescent detection of TH (red) and FITC-lectin IB4 (green). Sale bars: a) 50 μm, b) 30 μm, c) 50 μm and d) 20 μm.

### 3.3. NT3 activates lipolysis

The presence of TrkC in Wad suggest a functional role for this receptor. To clarify this role, we assayed its possible activity on lipolysis. Rat adipocytes isolated from WAT were incubated for 90 min without (control) or with NT3 (32 or 96 ng mL^-1^), the pan-Trk inhibitor K252a (0.2 μM) [35, 36] and NT3 + K252a. Glycerol production in presence of the different drugs was calculated as a percentage *vs* glycerol determined in the control. Because stimulation of β-adrenergic receptors reportedly increases lipolysis [37], we incubated the adipocytes in parallel with isoprenaline at 0.1, 1 or 10 μM. Incubation with NT3 (32 ng mL^-1^) induced a slight increase in glycerol release, similar to the one induced by the lower concentration of isoprenaline (Figure 3). A higher concentration of NT3 (96 ng mL^-1^) did not produce a higher response. The activity of NT3 was inhibited by pre-incubation with K252a (Figure 3). Albeit these result suggest that NT3 induces lipolysis in rat adipocytes through TrkC, the lipolytic activity of NT3 appears less efficient that the one induced by isoprenaline, suggesting that NT3 could be involved in other processes in AT.

**Figure 3.**
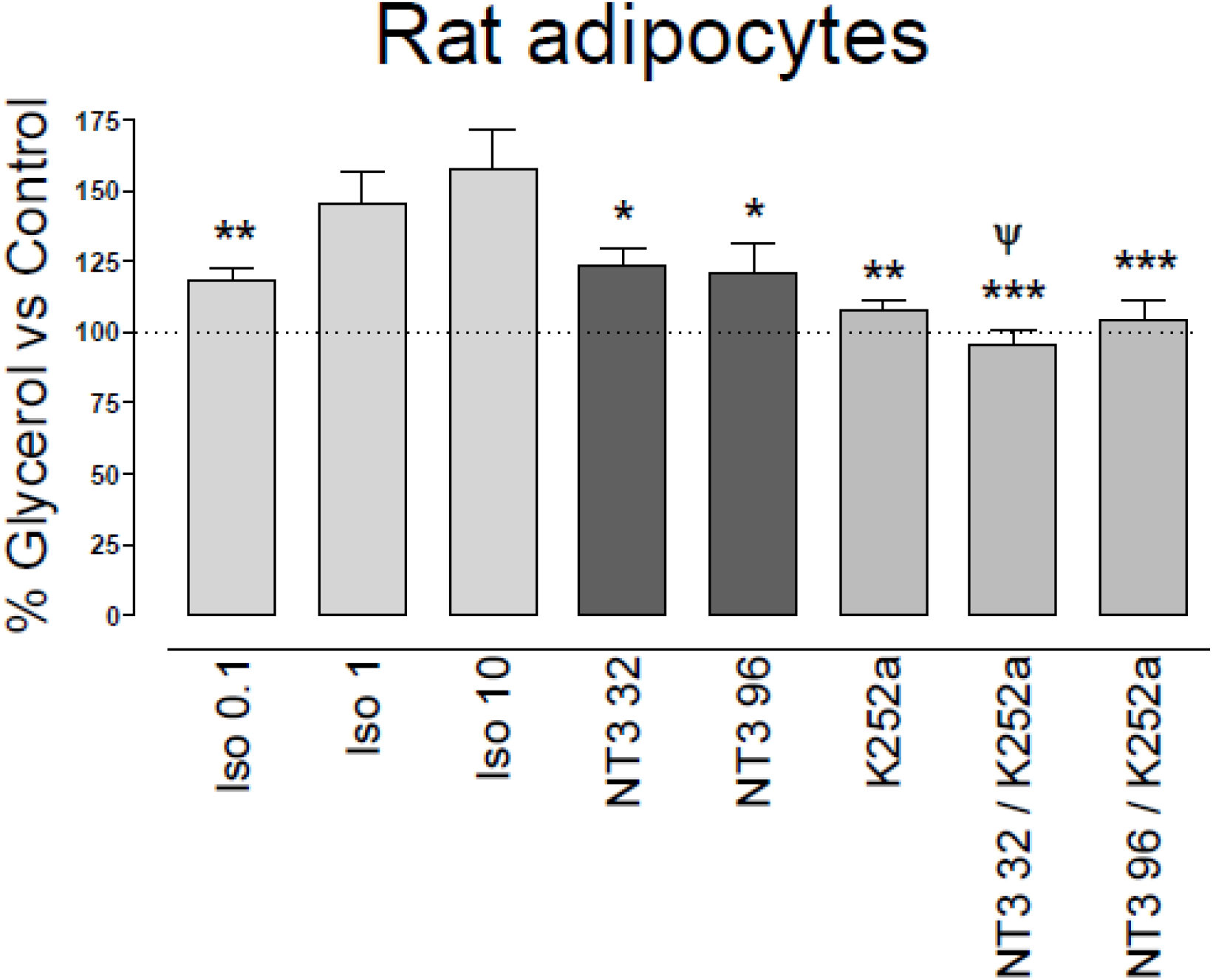
Lipolytic activity of NT3, through activation of TrkC, in isolated rat adipocytes. Isoprenaline (0.1 μM, 1 μM and 10 μM) or NT3 (32 and 96 ng mL^-1^) increases glycerol levels in adipocytes isolated from rat abdominal white adipose tissue. The Trk inhibitor K252a (0.2 μM) inhibits NT3-induced increase in glycerol levels. Data are expressed as a percentage *vs* control and are the mean± SEM of n= 15 experiments. * *P* < 0.05, ** *P* < 0.01, *** *P* < 0.001 vs isoprenaline 10 μM, ^ψ^ *P* < 0.05 vs NT3 32 ng mL^-1^, one-way ANOVA followed by Newman-Keuls test.

### 3.4. NT3 diminished human adipocyte size and increased UCP-1 expression

We have shown that NT3 modulates neural stem cell differentiation [24]. To analyze the possible activity of the NT3/TrkC pathway in human adipocyte differentiation, human pre-adipocytes were induced to differentiate into adipocytes for 15 days in absence (control) or presence of NT3 (32 ng mL^-1^), K252a (0.2 μM) and NT3 + K252a. Morphological analysis of the NT3 treated adipocytes showed a significant reduction of average size compared with untreated cells (Figure 4A and 4B). This reduction was not observed when adipocytes were cultured in presence of K252a, or in the combined presence of NT3 and K252a (Figure 4B). The diameter of lipid droplets was not affected by the treatment with NT3, although incubation with K252a induced an increase in the size of droplets not related to NT3 activity (Figure 4C). The lower adipocyte size observed in NT3 treated *vs* untreated cells was accompanied by an increased expression of UCP-1, a characteristic marker of brown and brite adipocytes, in human adipocytes differentiated in presence of NT3 (Figure 4D).

**Figure 4.**
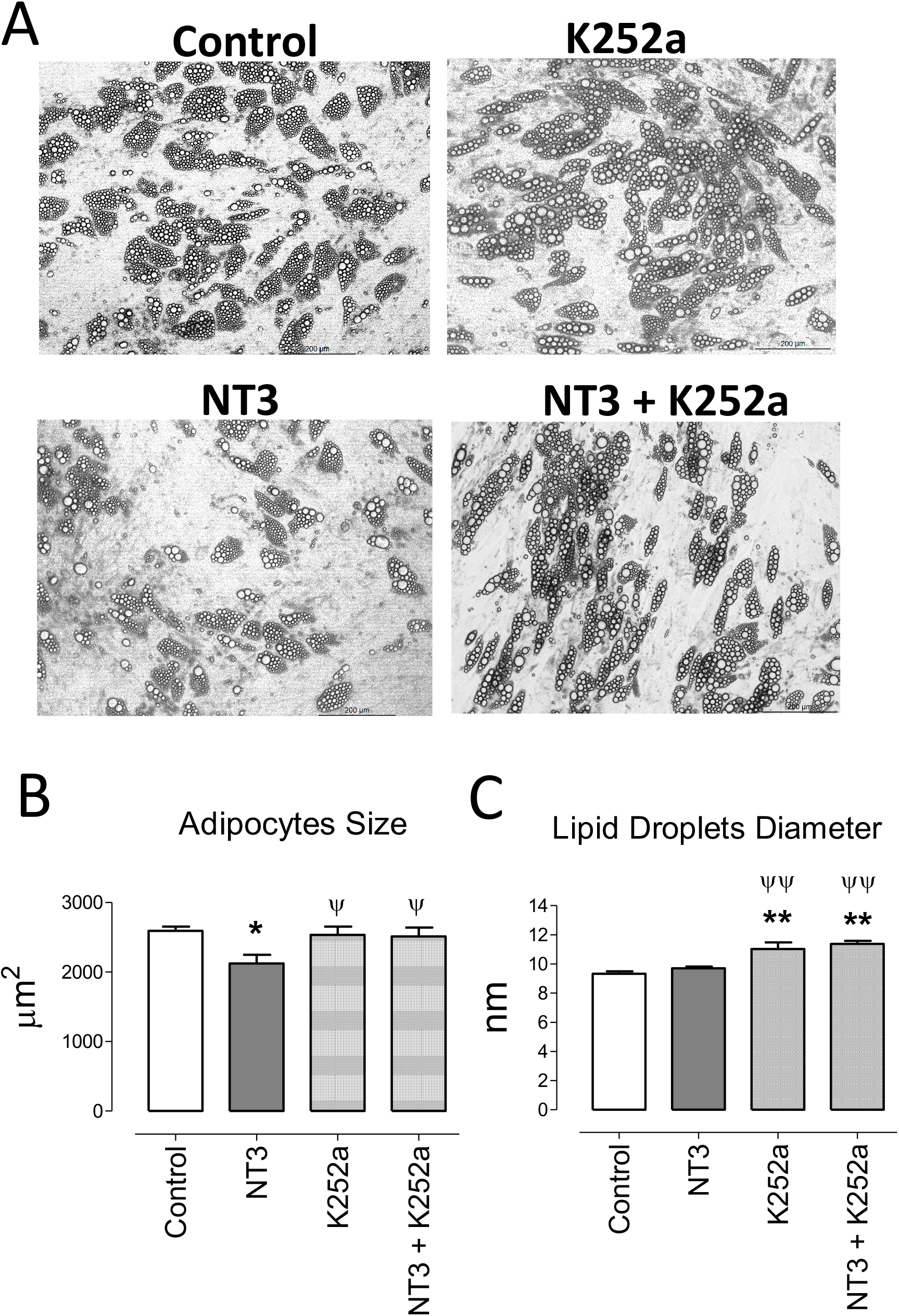
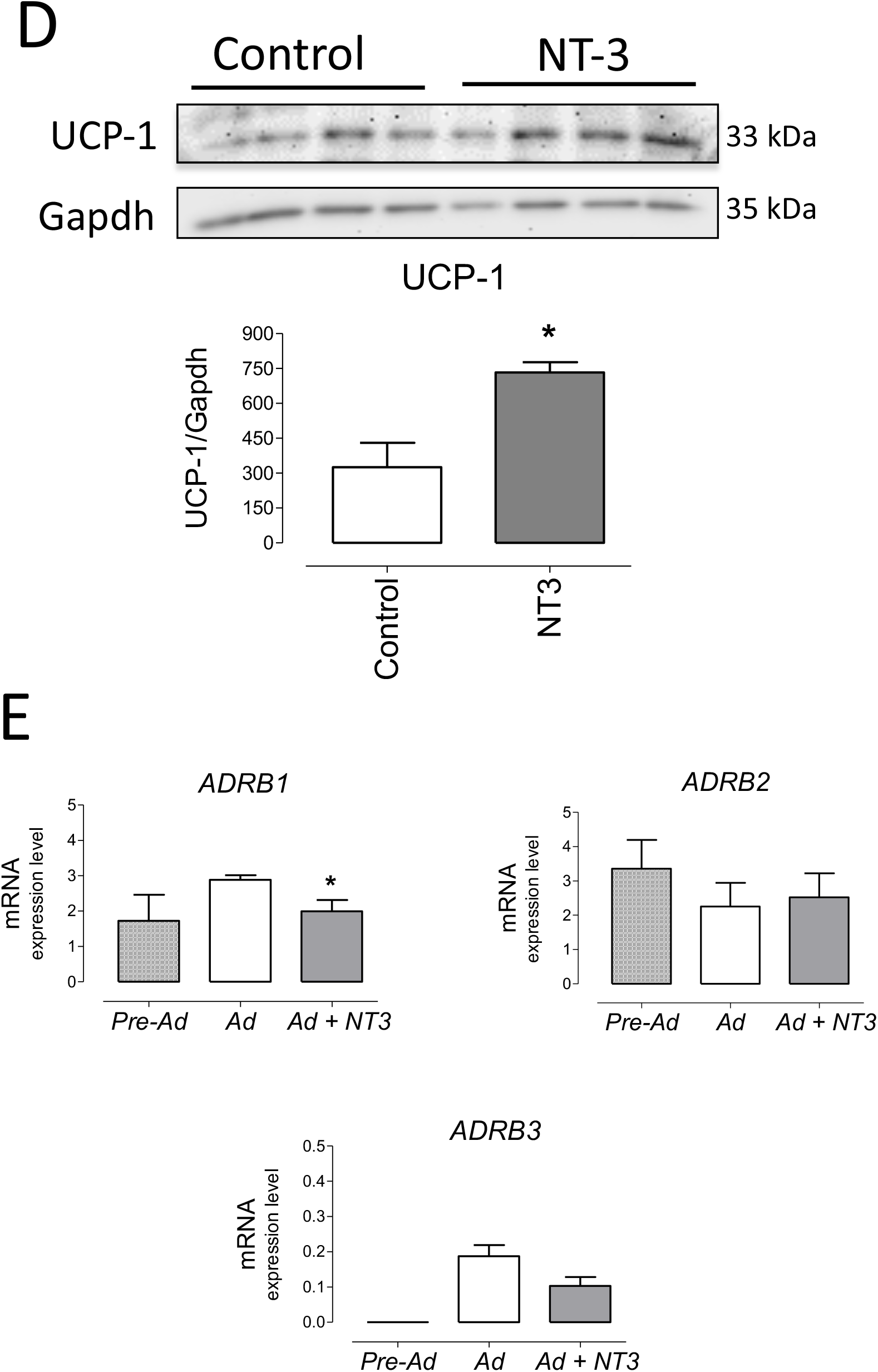
The NT3/TrkC pathway modulates human adipocyte differentiation giving mature adipocytes with lower size and higher UCP-1 expression. **(A)** Representative images of adipocytes differentiated from human pre-adipocytes in absence (control) or in presence of NT3 (32 ng mL^-1^), K252a (0.2 μM) and NT3 + K252a. Quantification of adipocytes size **(B)** and lipid droplets diameter **(C),** calculated from 5 different experiments performed in duplicate, 6 fields by plate, 10 cells and 25 lipid droplets by field, \**P* < 0.05, ** *P* < 0.01 vs control, ^ψ^*P* < 0.05, ^ψψ^ *P* < 0.01 vs NT3, one-way ANOVA followed by Newman-Keuls test **(D)** Representative immunoblot and quantification (bar diagram) of the expression of UCP-1 in human adipocytes differentiated in presence or not of NT3 (32 ng mL^-1^), \**P* < 0.05, Student’s *t* test. **(E)** Expression of *ADRB1, ADRB2 and ADRB3* in human preadipocytes (Pre-Ad) and adipocytes differentiated in absence (Ad) or presence (Ad + NT3) of NT3 (32 ng mL^-1^) *\*P* < 0.05 *vs* Ad, Student’s t test. Values of mRNA in bar diagrams are expressed as 2^-ΔCt^x10^4^ using *Gapdh* as a housekeeping gene. Data are mean± SEM of 5 different experiments.

The expression of *ADRB1, ADRB2* and *ADRB3* was determined in pre-adipocytes and human adipocytes differentiated in presence or absence of NT3 (32 ng mL^-1^). *ADRB3* was almost indetectable in preadipocytes whereas it was present in differentiated human adipocytes. mRNA levels for these genes did not significantly change (*ADRAB2*), tend to decrease (*ADRAB3*) or significantly decreases (*ADRAB1*) when NT3 was present during adipocyte differentiation (Figure 4E). In any case, the activity of NT3 in the differentiation of human adipocytes does not appear to be related to a greater expression of β adrenoceptors.

### 3.5.

Genetically engineered mice with deficient expression of NT3 exhibit adipocytes with higher size and lower UCP-1 expression

To functionally analyze the role of NT3 in AT *in vivo* we used *Ntf^3+/lacZ^ mice* (referred to as NT3^+/-^), which express reduced levels of NT3 [11] and *Ntf3^flox1/flox2^;Tie2-Cre^+/0^* mice (referred to as eNT3-) that selectively prevents NT3 expression in endothelial cells [24], compared to their controls *Ntf3^+/+^* (NT3+/+) and *Ntf3^flox1/flox2^;Tie2-Cre^0/0^* (eNT3+) mice, respectively. We first confirmed the reduced expression of *Ntf3* in genetically engineered mice and analyze possible changes in the expression of the other genes studied. As expected, a lower expression of *Ntf3* was found in WAT and BAT from NT3^+/-^ mice and eNT3-mice, compared to their littermates (Figures 5A and 5B). Not significant differences were found in the expression of *NTrk3*, nor in *Adrb1, Adrb2* and *Adrb3* expression in WAT and BAT from NT3^+/+^ and NT3^+/-^ mice (Figure 5A). However, decreased expression of *Ntrk3* and *Adrb3* was found in WAT and BAT from eNT3- *vs* eNT3+ mice (Figure 5B). No significant differences were observed in the body weight, fat depots, glycemia and plasmatic lipid profile between NT3^+/+^ and NT3^+/-^ mice nor between eNT3- and eNT3+ mice (Table 2). Interestingly, an increase in the average size of adipocytes was found in WAT and BAT from NT3^+/-^ *vs* NT3^+/+^ (Figure 5A) and from mice without endothelial expression of NT3 (eNT3-) (Figure 5B). Moreover, a decrease of UCP-1expression was observed in BAT samples from eNT3-mice compared to eNT3 mice (Figure 5C). Altogether, these data indicate that endothelial NT3 modulates size and UCP-1 expression in mouse adipocytes not only *in vitro* but also in *vivo* conditions.

**Figure 5.**
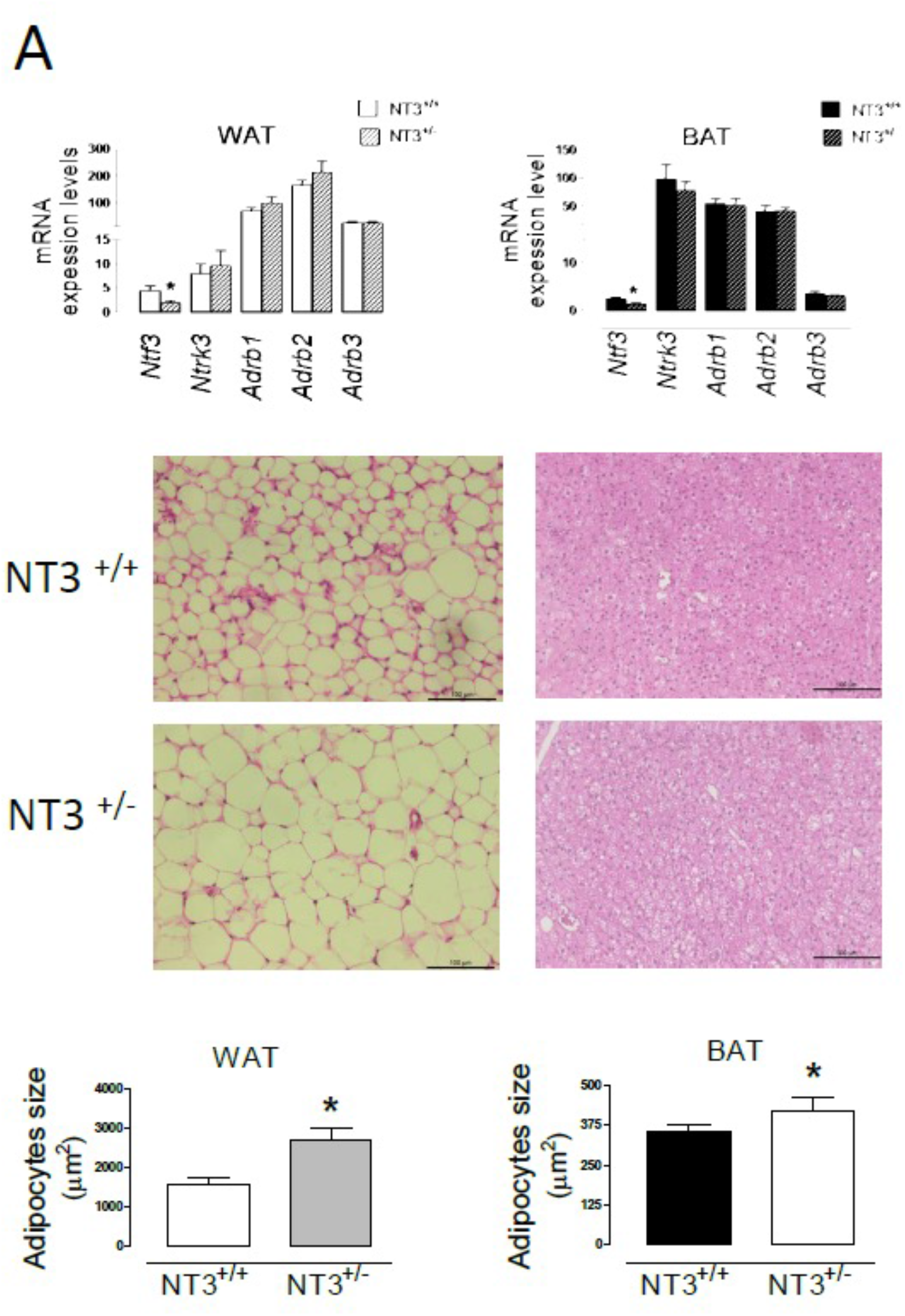
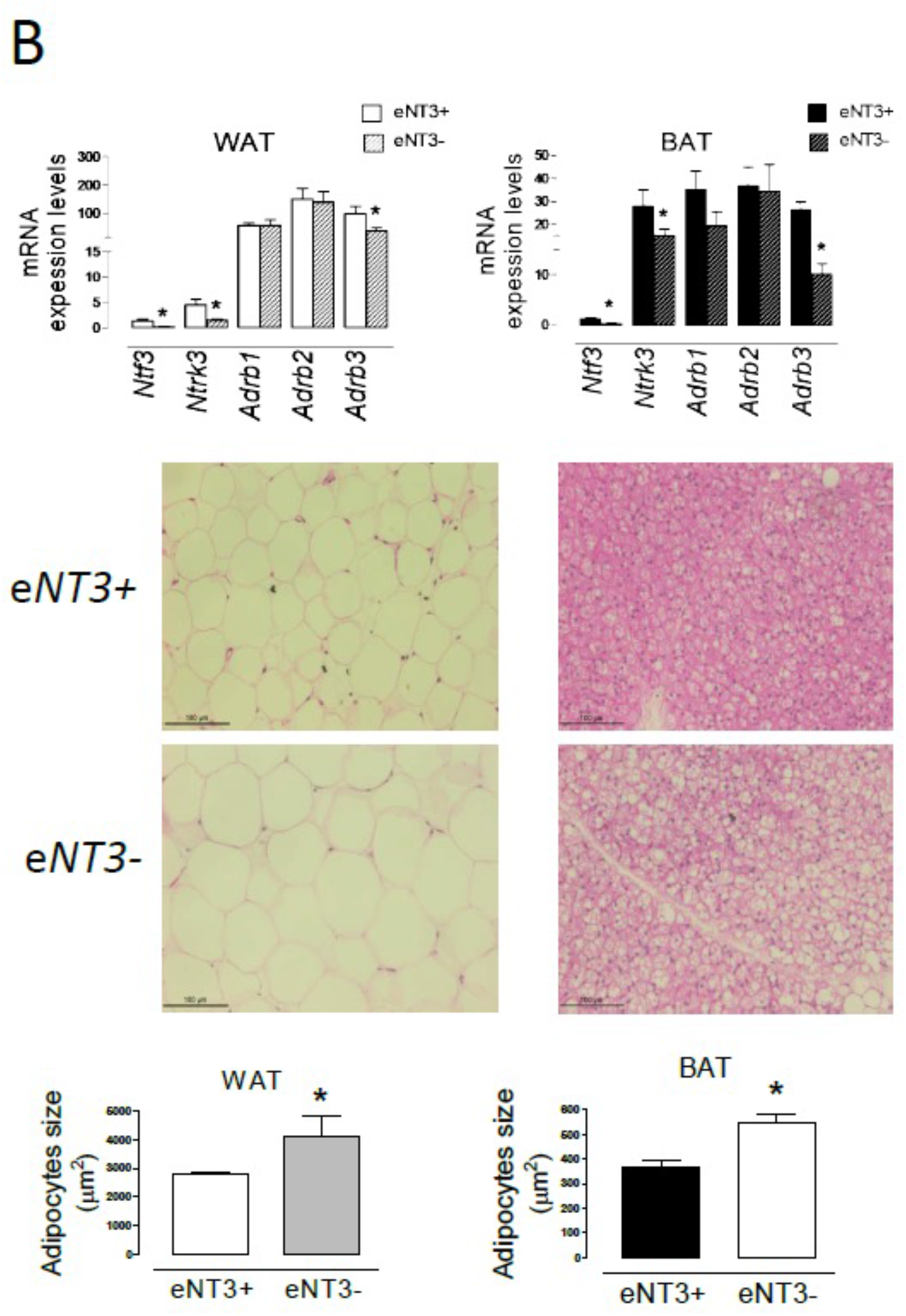
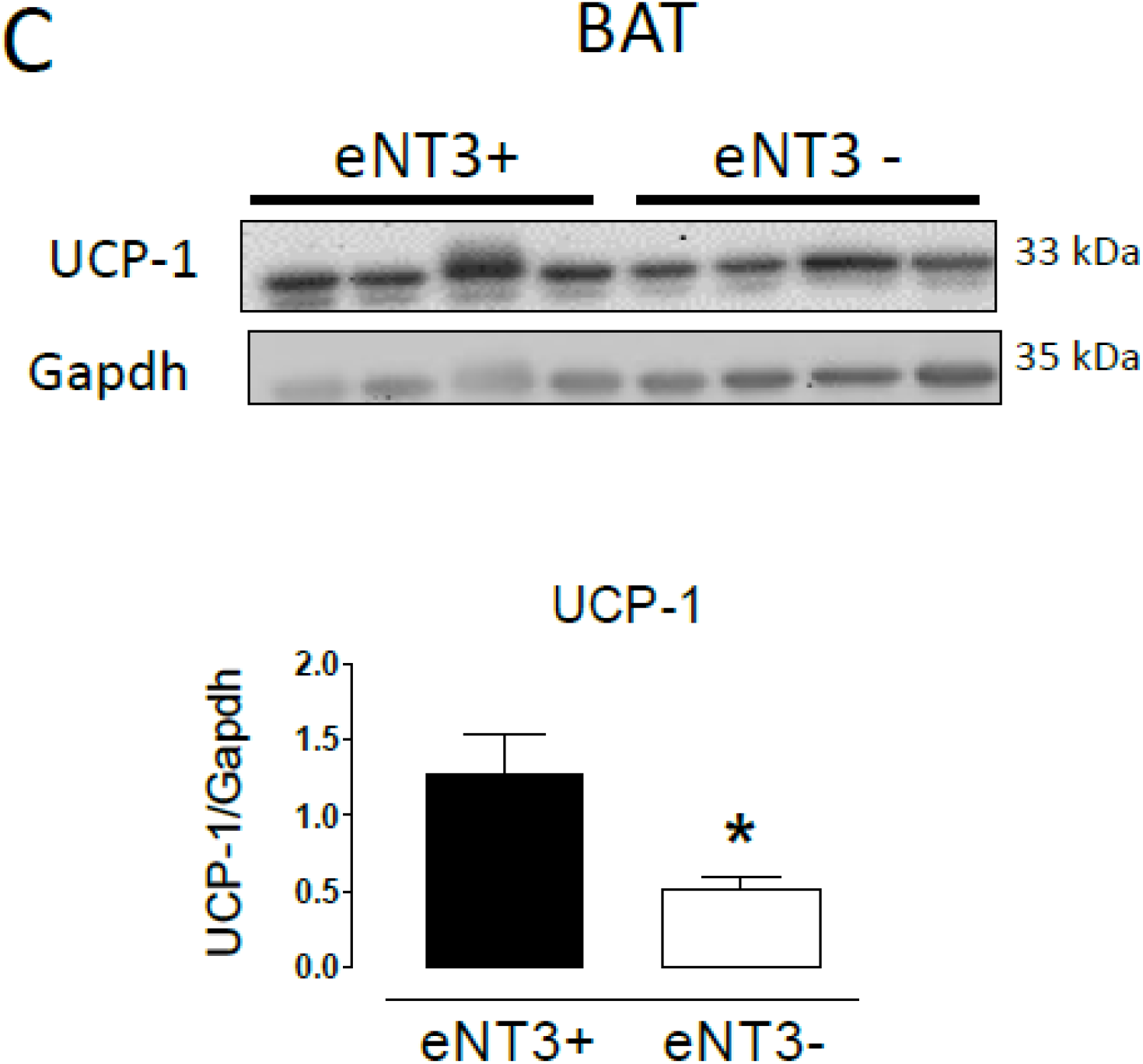
Deficient expression of NT3 is related to higher adipocyte size and lower UCP-1 expression in mice adipose tissue. mRNA levels of genes (*Ntf3, Ntrk3, Adrb1, Adrb2, Adrb3*), paraffin-embedded sections of tissues stained with hematoxylin/eosin and quantification of the adipocyte size, in WAT and BAT from male NT3 ^+/+^ and NT3 ^+/-^ mice **(A)** and male *eNT3*- and their control littermates (eNT3+) **(B), (C)** Representative immunoblot and quantification of protein expression of UCP-1 in BAT from eNT3- and eNT3+ mice. Data are mean± SEM of 4-6 different animals per group. \**P* < 0.05 *vs* their littermates, Student’s t test

**Table 2.** Analysis of body weight, fat depots and plasma biochemical markers in genetically engineered mice with deficient expression of NT3: *Ntf3^+/lacZneo^* mice (NT3^+/-^) and their littermates (NT3^+/+^) and mice without endothelial expression of NT3 (*Ntf3^flox1/flox2^;Tie2-Cre^+/0^*: eNT3-) and their littermates (eNT3+)

|  | **NT3<sup>+/+</sup>**<br>**(n=8)** | **NT3<sup>+/-</sup>**<br>**(n=6)** | **eNT3+**<br>**(n=11)** | **eNT3-**<br>**(n=11)** |
| --- | --- | --- | --- | --- |
| **Age, weeks** | 35.8 ± 0.4 | 38.5 ± 2.3 | 57.1 ± 5.1 | 53.3 ± 3.0 |
| **Body weight, g** | 38.1 ± 1.7 | 37.2 ± 2.7 | 34.3 ± 0.4 | 31.7 ± 1.2 |
| **WAT/Body weight, g** | 1.57 ± 0.51 | 1.64 ± 0.56 | 2.52 ± 0.21 | 2.17 ± 0.57 |
| **KAT/Body weight, g** | 0.08 ± 0.05 | 0.04 ± 0.03 | 0.11 ± 0.03 | 0.15 ± 0.07 |
| **BAT/Body weight, g** | 0.46 ± 0.06 | 0.42 ± 0.07 | 0.27 ± 0.05 | 0.38 ± 0.09 |
| **Glucose, mg/dL** | 139.2 ± 15.5 | 160.7 ± 23.4 | 180.0 ± 13.86 | 174.2 ± 13.2 |
| **Total cholesterol, mg/dL** | 133.4 ± 4.2 | 129.5 ± 7.1 | 98.3 ± 7.1 | 86.6 ± 4.7 |
| **HDL-cholesterol, mg/dL** | 70.06 ± 4.3 | 72.7 ± 3.9 | 40.9 ± 5.1 | 38.0 ± 3.6 |
| **LDL-cholesterol, mg/dL** | 49.1 ± 4.3 | 34.7 ± 5.0 | 50.3 ± 6.4 | 38.0 ± 2.2 |
| **Triglycerides, mg/dL** | 149.4 ± 24.3 | 154.0 ± 39.0 | 68.3 ± 5.9 | 78.4 ± 8.1 |

## 4. DISCUSSION

The major findings of present study are: i) both NT3 and its cognate receptor TrkC are expressed in human and rodent AT; ii) NT3, but not TrkC, expression decreases with ageing. iii) NT3 is mainly expressed in vessels irrigating adipose tissue, whereas TrkC expression is similar in isolated adipocytes and the entire tissue. iv)human adipocytes differentiated in the presence of NT3 exhibit decreased size and increased UCP-1 expression; v) mice with reduced expression of NT3 (NT3^+/-^), or with prevented endothelial NT3 expression (eNT3-), exhibit larger adipocytes, in WAT and BAT, and lower UCP-1 expression in BAT. Collectively, these observations support the notion that NT3, mainly released by blood vessels irrigating the AT, exerts an important regulatory activity on adipocyte differentiation and physiology through activation of TrkC. To our knowledge, this is the first evidence of the presence of a functional NT3/TrkC pathway in AT.

### 4.1. The NT3/ TrkC pathway is present in human and rodent adipose tissue

WAT controls whole-body energetics and fuel metabolism. In contrast, BAT dissipates energy and generates heat by thermogenesis, through UCP-1-mediated mitochondrial uncoupling. WAT shows fewer nerves and lower number of blood vessels than BAT, which is richely vascularised and innervated with noradrenergic fibers, in direct contact with the brown adipocytes [37]. Sympathetic innervation and the adrenoceptor subtypes present contribute to multiple aspects in the regulation of adipocyte function including lipogenesis, lipolysis, glucose metabolism, thermogenesis, and the secretion of hormones. In WAT, the sympathetic nervous system through β-adrenoceptor activation, is the principal initiator of lipolysis [38–41]. In BAT, β-adrenergic activation leads to increased expression and activation of UCP1 in the inner mitochondrial membrane, which collapses the proton gradient that would normally drive ATP synthesis and energy storage, increasing mitochondrial respiration and thermogenesis [28].

In rodents, all three β-adrenoceptor subtypes are expressed in subcutaneous and visceral adipose depots with the β3-adrenoceptor being the main responsible for β-adrenoceptor-mediated lipolysis in mature white adipocytes [42]. In human white adipocytes, expression of β3-adrenoceptors is not as relevant as the other subtypes [42]. Our results confirm the abundant expression of *Adrb3* in rat ATs, or *ADRB2* and *ADRB1*, with minor levels of *ADRB3*, in hAT. However, according to present results, a preeminent role in the modulation of human body weight could be assumed for the β3-subtype, since its expression in hAT inversely correlates to BMI but no relationship was observed as regards of β1- or β2-adrenoceptors. In this way, our results coincide to previous observations about a close relationship between β3-adrenoceptors and long-term changes in body weight in adulthood [43].

Interestingly, *ADRB2*, the mainly expressed subtype in hAT, did not correlate to any of the clinical variables, but its expression was significantly lower in patients with a sedentary style of life. This observation agrees with previous reports which indicates that β-adrenoceptor modulation of thermogenic responsiveness is augmented in regularly exercising compared with sedentary adults [44]

NT3 modulates development and survival of the adrenergic neurons but its presence, function and relationship to adrenergic activity in AT had not been previously analyzed. Present results consistently indicated that NT3 and TrkC are both present in hAT and in different rodent ATs as WAT, BAT, VAT and KAT. It is remarkable that *Ntf3* or *Ntrk3* expression in AT is comparable, or even higher as *Ntrk3* in BAT, to that found in cerebral cortex. Furthermore, expression of TrkC in human or rat ATs is similar to that of β-adrenoceptors, essential regulators of the AT function, suggesting a potential role for this receptor in AT.

Previous papers have described controversial but significant correlations between BMI or cholesterol (total or LDL-cholesterol) and plasmatic levels of NT3 in morbidly obese or alcoholic patients [26,27]. Our results did not show any significant correlation between *NTF3* in hAT, and BMI or lipid profile, although the sample size does not allow us to obtain definitive conclusions. However, in hAT, and inverse correlation was found between *NTF3* and *ADRB3* levels with age. Similarly, a decreased expression of *Ntf3* and *Adrb3* was observed in aged *vs* young rats. The possible relationship between the reduced expression of β3-adrenoceptor and NT3 in AT, and the previously described age-related changes in lipid metabolism [45] needs further investigation.

It has been described that the vascular bed, and more specifically the endothelial cells, produce a large amount of NT3 that facilitates the growth of axons and the development of adrenergic innervation [24, 25]. Confirming these observations, our results show, in AT, the proximity of the sympathetic fibers to vessels, the main source of NT3. This characteristic location could suggest indirect neurogenal effects of NT3 in lipid metabolism but also, that *Ntf3* detected in AT could have a vascular origin. In fact, our samples of AT could include microvessels and we can suppose that the *Ntf3* found in AT has a vascular origin. For this reason, *Ntf3* levels in isolated adipocytes were significantly lower than in the entire tissue. However, there are arguments that do not permit us to exclude the expression of NT3 in adipocytes. First, Ntf3 was detected in isolated adipocytes. Second, the observation that, in spite of the rich vascularization and innervation characteristic of BAT [28, 38], similar or higher levels of *Ntf3* were found in WAT and KAT vs BAT. But, independently of NT3 origin, the presence of its cognate receptor TrkC could justify the relevance of the pathway in these cells. In this way, mRNA levels of *Ntrk3* in adipocytes isolated from rat WAT mimic the TrkC expression pattern of the entire tissue, and this observation suggests that NT3 mainly from vascular origin, has a role not only as a regulator of neuronal growth in the AT, but better, as a paracrine modulator of the adipocyte function. Therefore, our next step was to investigate this role.

### 4.2. NT3 stimulates lipolysis, decreases adipocyte size and increases UCP-1-expression

The presence of TrkC in adipocytes and the stimulation of lipolysis by NT3 on isolated adipocytes suggest a direct role for the NT3/TrkC pathway in AT beyond the modulation of sympathetic innervation. Addition of NT3 slightly increased glycerol accumulation, an increase that was blocked by K252a, a non-selective Trk-receptor antagonist [35,36]. However, the magnitude of the NT3 response, indicates that this is not a remarkable function compared to β-adrenoceptor activation mediated by isoprenaline. Therefore, these observations lead us to suppose that the presence of NT3 and TrkC in AT could have another purpose.

We have previously shown that NT3 modulates neural stem cell differentiation in the adult mouse brain [24]. On this basis, we hypothesize that NT3 present in AT could also modulate adipocyte differentiation. Consistent with our proposal, human preadipocytes differentiated in presence of NT3 produced smaller mature adipocytes, an effect that is mediated by TrkC, as K252a blocks it. The decrease in adipocyte size induced by NT3 was not accompanied by changes in lipid droplet diameter. Rather, K252a promotes an increase in droplet diameter not related to NT3 activity. As K252a is a pan-Trk inhibitor, its activity on the other Trk receptor subtypes could explain this result but needs further investigation.

More interestingly, human mature adipocytes differentiated in the presence of NT3 exhibit a higher expression of UCP-1, the major determinant of non-shivering thermogenesis and a characteristic marker of mature brown and brite adipocytes. The ability to drive uncoupled respiration through UCP-1 activation represents a possibility in the treatment of metabolic disorders by expending excess energy as dissipated heat [42, 46]. The ability to induce an increase in the expression of UCP-1 is the first consequence of the activation of β-adrenoceptors, especially the β3 subtype. However, present results show that NT3 activity on UCP-1 expression does not depends on increased β-adrenoceptors population, since similar mRNA levels of ADRB2 and ADRB3 and lower levels of ADRB1, were found in adipocytes differentiated in presence or absence of NT3. In fact, as has been previously described [42], human adipocyte cultures display negligible expression of ADRB3, even after differentiation in the presence of highly adipogenic stimulus.

Whether the change in UCP-1, observed by NT3 treatment of cultured human adipocytes, could be translated to *in vivo* AT remains debatable, due to technical limitations. In fact, differentiation of pre-adipocytes into adipocytes follows a protocol that induces lipid accumulation and the formation of adipocytes with multilocular appearance, regardless of the tissue origin, and devoid of tissue-specific regulatory mechanisms that drive cell differentiation and metabolic adaptative responses [47]. Therefore, adipocytes differentiated by this procedure exhibit a phenotype characteristic of brite or beige adipocytes, with a mixture of small and large lipid droplets and UCP-1 and β3-adrenoceptor expression [42].

For this reason, our next objective was to corroborate these observations *in vivo*, using two models of NT3-deficient mice: heterozygous *Ntf3* mice (*NT3*^+/-^), and mice genetically modified to eliminate NT3 expression in endothelial cells (*e*NT3-). The results obtained showed an increased size of adipocytes in WAT and BAT obtained from both models and, as occurred in human adipocytes, this activity could not be related to changes in β-adrenoceptors expression since it was not altered in NT3^+/-^ mice.

Endothelial cells are an important source of NT3 [24, 25 and present results]. Consistent with this, abrogating endothelial production of NT3 results in a phenotype in AT, that resembles that of heterozygous *Ntf3* mice, with more voluminous adipocytes both in WAT and BAT. Importantly, expression of the UCP-1 was decreased in *e*NT3- mice, as occurred in human adipocytes differentiated in absence of NT3.

It is well known that β-adrenoceptors, and specially the β3 subtype, control the transcription of nuclear factors that increase mitochondrial biogenesis and UCP-1 expression [37, 41, 42, 48, 49]. However, the generation of β3 knockouts has not resulted remarkable changes in AT [50] by functional redundancy between the three β-adrenoceptor subtypes co-expressed in adipocytes [42, 51]. Only the mice that lack all three β-adrenoceptors exhibit a BAT with large cells and lower levels of UCP-1 [52], a phenotype similar to that observed in mice with partial *Ntf3* expression or lack of endothelial *Ntf3*. Therefore, the importance of the NT3/TrkC pathway in the modulation of adipocyte differentiation and activity could be comparable to β-adrenergic activity. Furthermore, the location of the sympathetic fibers that innervate the AT in the proximity of the vessels (the main source of NT3), suggests that endothelial NT3 could posses an additional activity, regulating the growth and function of sympathetic terminals.

### 4.3. CONCLUSIONS

Until now, pharmacological modulation of β-adrenoceptors, especially the β3 subtype, had been a promising strategy to the treatment of obesity, and considerable efforts had been made to develop β3-adrenoceptor agonists. Unfortunately, this approach has not yet been successful [53, 54]. Different reasons could explain the divergence between humans and rodents. While β3-adrenoceptors play a major role in rodent AT, adult humans have white AT in which β3-adrenoceptors are poorly expressed and play only a minor role in lipolysis [54, 55]. In spite of this, our results suggest a relevant role for this subtype in the control of obesity, since the β3-, but not β1-nor β2-adrenoceptor expression in hAT, inversely correlates to body mass index. Interestingly, a decreased expression of the predominant subtype, the β2, was related to a human sedentary way of life. Another question is the existence of undefined differences between rodent and human receptor physiology which implies a poor activity of agonists at the human *vs* rodent β3-adrenoceptor [42]. Furthermore, β3-adrenoceptor expression and function decrease with age (present results), obesity or diabetes [42], then, it is plausible that patients who would benefit most from activation of β3-adrenoceptors by selective agonists may be less able to respond them. In this way, our results indicate that NT3, also decreases with age in human and rat AT but its specific receptor TrkC did not. As stimulation of TrkC by NT3 mimics the effects of β-adrenoceptors on UCP1 expression and adipocyte size in human and rodents, a drug selectively acting on TrkC, abundantly expressed in WAT and BAT, could be a promising pharmacological target in the management of obesity.

In summary, our study is the first to describe the presence of NT3/TrkC pattern on human and rodents AT, and to demonstrate a direct action of this pathway on the regulation of adipocyte differentiation and UCP-1 expression, *in vitro* and *in vivo*. This evidence opens new perspectives for future investigations about the role of the NT3/TrkC pathway in cardiometabolic disorders and the search of new drugs acting on this target.

## ACKNOWLEDGEMENTS

We are grateful to M Carmen Iglesias-Osma and M Jose Barrado (Departamento de Farmacología, Facultad de Medicina, Universidad de Salamanca, Spain) for helpful advice with the development of the lipolysis experiments. The authors thank the technical support of the Microscopy Service and the Animal Production Service from the Servicio Central de Apoyo a la Investigación Experimental (SCSIE), Universidad de Valencia.

## STATEMENT OF ETHICS

This research conforms with the principles outlined in theWorld Medical Association Declaration of Helsinki and was performed with the approval of the Ethic Committee of the Hospital Clínico Universitario San Carlos, Madrid, Spain with registry number 19/011. Animal handling and all experimental procedures were carried out according to European Union 2010/63/UE and Spanish RD-53/2013 guidelines, following protocols approved by the institutional ethics committee of the University of Valencia.

## AUTHOR CONTRIBUTIONS

MB, JTR, IF and PD contributed to conception and design of the study; MGA and TT obtained human samples and clinical data; MB, FM, PGLl, AZ, MAN, ACR and SSP performed experiments; MB, FM PGLl, DI and PD organized the database and analyzed data; PD is the main writer of the manuscript. All authors contributed to manuscript revision, read and approved the submitted version.

